# Latitudinal gradients in air density create “invisible topography” at sea level affecting animal flight costs

**DOI:** 10.1101/2024.08.21.608961

**Authors:** Emily L.C. Shepard, Baptiste Garde, Krishnamoorthy Krishnan, Adam Fell, Vikash Tatayah, Carl G. Jones, Nik C. Cole, Emmanouil Lempidakis

## Abstract

Regional patterns in wind underpin the low-cost migratory flyways of billions of birds and insects^1-3^, but how large-scale changes in temperature affect flight is unknown. Flight costs should increase with rising temperatures, because lift decreases as density decreases, whereas weight remains unchanged. The effects of air density on flight costs are well-established in the context of high-altitude movements and migration^4-7^. Here, we examine the impact of air density on low-flying birds, in relation to seasonal, regional and global changes in temperature. Using multi-sensor loggers, we find that air density was the most important predictor of wingbeat frequency in red-tailed tropicbirds (*Phaethon rubricauda*) breeding year-round in Mauritius. Lower air densities in the Austral summer were associated with a small but significant increase in mean wingbeat frequency, which translated to an estimated 1-2% increase in flight costs. The variation in flight costs increased by an order of magnitude when considered in space, rather than time, with flight costs varying by ≥ 10 % across the tropicbird’s range. Changes in air density can therefore be an important determinant of flight costs even when birds are operating close to sea-level. Indeed, mapping air density at sea-level revealed that global temperature gradients cause effective altitude to vary by >2 km when considered as seasonal averages. This “invisible topography” at sea-level could have influenced the biogeography of flight morphologies and life-history traits.

## RESULTS

Tropical seabirds represent model systems with which to examine how flight effort is affected by the physical flight environment as some species breed year-round, and hence experience a range of weather conditions^8,9^. We used high-frequency movement data to quantify flight effort in red-tailed tropicbirds (*Phaethon rubricauda*) breeding in the Austral summer and winter on Round Island, Mauritius. Specifically, we quantified how wind and air density affected wingbeat frequency (which is proportional to power output ^10^), controlling for the effect of prey loading on flight costs by quantifying prey encounter rates.

Data were recorded from 55 red-tailed tropicbirds undertaking 76 foraging trips; 31 in the Austral summer (February-March) and 45 in the Austral winter (September-October). There was no evidence for seasonal variation in body mass (Student’s t-test: t = 0.282, p-value = 0.779) or wing area between seasons (Student’s t-test: t = -0.773, p-value = 0.446). Birds covered an average of 100.1 km in 5.4 hours during their foraging trips and distance remained consistent between the two seasons (LME: std. error = 2.085, t-value = 1.343, p = 0.185) (Fig. 1). Prey encounter rate did not vary significantly between seasons (Wilcoxon test: W = 857, p-value = 0.093), with one pursuit being recorded every 8.0 km during the summer, and one every 10.8 km in the winter. Foraging efficiency (the number of pursuits divided by the total number of wingbeats per flight) also appeared constant across seasons (LME: t-value = -1.342, p = 0.185, Table S1). The lack of a seasonal difference in foraging success was surprising given that breeding activity increases 5-fold from summer to winter in Mauritius (Fig. S1), but our metrics of foraging success are based solely on an increase in body mass and ignore potential differences in the energy density of prey.

**Figure 1.**
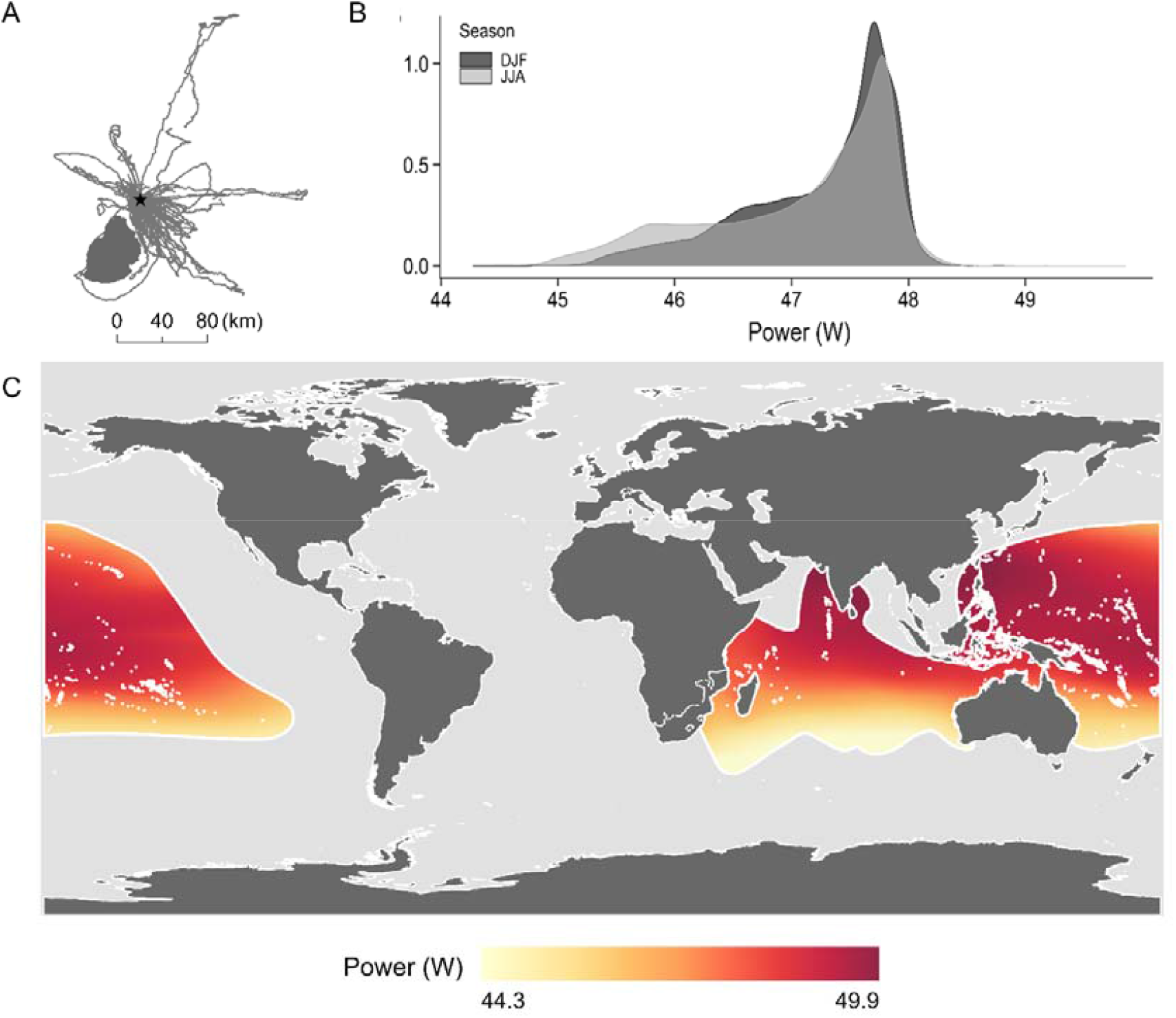
(a) Foraging movements of red-tailed tropicbirds from Round Island (indicated with a black star, lying to the north of mainland Mauritius). (b) The frequency of flight power values estimated across the red-tailed tropicbird distribution during the Austral summer (December, January, February, DJF) and winter (June, July, August, JJA). (c) Spatial variation in flight power as a function of regional variation in air density within the red-tailed tropicbird range. The power required to fly was estimated using apft^17^, using the morphological measurements of a red-tailed tropicbird of average mass from Round Island flying at its maximum range speed (V_mr_), in varying air densities. The breeding distribution is taken from BirdLife International (http://datazone.birdlife.org/species/requestdis). Monthly mean values of air density were estimated from ERA5 for 2001-2020 (https://cds.climate.copernicus.eu/cdsapp#!/dataset/reanalysis-era5-single-levels-monthly-means?tab=overview)

Average air temperatures varied by 8.4 °C across all flights and the mean temperature in the summer was 4.1 °C higher (Wilcoxon test: W = 378, p < 0.001). Air density varied by 0.06 kg m^-3^ across flights and was 0.03 kg m^-3^ lower on average in the summer (Student’s t-test:, t = -19.542, p < 0.001, Table S1, Fig. S1). Flight altitudes varied slightly between seasons, being 13.5 m higher on average during the summer (LME: estimate = -13.542, std. error = 5.050, t-value = -2.681, p = 0.010, Table S1).

Airspeed was predicted by wind and air density (raw estimate = -35.3, R^2^_m_ = 0.18, R^2^_c_ = 0.25, Table S2). Birds increased their airspeed with decreasing air density and increasing altitude, in line with theoretical predictions (Table S2). There was no collinearity between altitude and density, showing that air density varied between seasons and independently from flight altitude. There was no collinearity between wind speed, headwind and crosswind component, but headwind component and wind speed had an effect of similar magnitude (scaled estimates: 0.82 for headwind and 1.03 for wind speed), suggesting that the full effect of the wind was not captured by our estimates of the headwind component alone. Airspeeds did not vary between the two seasons (LME: std. error = 0.373, t-value = -1.329, p = 0.188, Table S1), consistent with the lack of a seasonal difference in mean wind speed but slightly at odds with the seasonal difference in air density.

A simple comparison of wingbeat frequency in level flapping flight showed a difference of 0.1 Hz between seasons, with flight being more costly in the summer (LME: estimate = -0.096, std. error = 0.030, t-value = -3.194, p = 0.002). Our LME model showed that air density was the most important predictor of wingbeat frequency across seasons (scaled estimate = -0.084, LME: raw estimate = -5.392, p = 0.004, Table S3). Flight altitude was also significant (scaled estimate = -0.061, p < 0.001).

We did not find any effect of airspeed on wingbeat frequency (p = 0.226), although airspeed did have a positive effect on the slope of the relationship between air density and wingbeat frequency (Raw estimate = 0.266, p = 0.014), such that the effect of air density on wingbeat frequency was reduced at high speed. Wingbeat frequency did increase with the number of prey pursuits (Scaled estimate = 0.031, p < 0.001), but the slope of the relationship between number of pursuits and wingbeat frequency did not vary between seasons (Table S3). Overall, the variation explained by the fixed effects was low compared to the random effect of individual (R^2^m = 0.13 and R^2^c = 0.77 respectively).

## DISCUSSION

Air density has a fundamental impact on the power required to fly^11^, because lift decreases as density decreases, whereas weight remains unchanged. Birds flying at lower air densities must increase their wingbeat frequency and associated lift production to achieve the same airspeed^11,12^, as well as fly faster to generate appropriate lift in rarefied air, all of which requires higher power. To date, the effect of air density on flight costs has only been considered when birds make substantial changes in flight height, as recorded for birds migrating over the Himalayas (≤ 7,290 m, Bishop *et al*. ^6^) the Sahara^13^, or for hummingbirds occurring across marked altitudinal ranges^14^. We demonstrate that temperature-driven changes in density can affect flight effort independently of flight altitude. Indeed, air density was the most important predictor of wingbeat frequency across seasons, with flight costs increasing in the Austral summer. The seasonal difference in air density (0.03 kg m^-3^) is equivalent to birds flying ∼ 250 m higher in the summer on average (assuming standard atmospheric conditions). This is much greater than the 13.5 m difference in mean flight height between seasons.

The increment in wingbeat frequency that we observed from winter to summer (which appears to be mostly attributable to the changing air density), is relatively small (0.1 Hz). However, it is twice what is predicted by Pennycuick’s theoretical framework (Flight version 1.24) for the mean seasonal air densities in our study (taking wing measurements from our study and wingbeat frequency at the minimum power speed as a reference speed^10^). It is possible that the energetic consequences of increases in wingbeat frequency may be mitigated by a reduction in wingbeat amplitude We were unable to assess this, as small changes in tag attachment between seasons influenced the amplitude of the acceleration signal^15^, but we note that hummingbirds increase their wingbeat amplitude with decreasing density^14^. Alternatively, low air densities could be more costly than predicted by theoretical models. Indeed, two previous studies also found that wingbeat frequency varied to a greater extent with altitude than predicted by theory for a range of migratory species^11,16^. Furthermore, both wingbeat frequency and the metabolic costs of flight increased more rapidly than anticipated due to declining air density in bar-headed geese (*Anser indicus*) migrating over the Himalayas^6^.

We can model the extent to which seasonally changing temperatures affect tropicbird flight costs, first using a classic aeronautical model^10^, and then modifying this with the results from the bar-headed geese^6^. We first use afpt: Tools for Modelling of Animal Flight Performance^17^ to predict the chemical power required for a tropicbird to fly at its maximum range speed across a range of air densities (taking a bird with average mass for our study). This provided the intercept of the relationship between power and density. We then took the slope of this relationship as (i) *P*_*m*_ ∝ *ρ*^−0.54^, as specified by Pennycuick ^10^ and (ii) *P*_*m*_ ∝ *ρ*^−0.91^, as proposed by Bishop *et al*. ^6^ following their empirical results from bar-headed geese, and used both to predict changes in flight power between mean seasonal air densities in Mauritius. This suggests that the mean power requirements of red-tailed tropicbirds vary by 1.4 - 2.4% between seasons, according to the proportionalities given by the two frameworks respectively.

While the seasonal change in mean power costs in one location is predicted to be low, decreases in air density will provide an additional cost to breeding in high temperatures beyond known stressors e.g. heat stress^18^ and declining ocean productivity^19^. This highlights a key difference between the effects of wind and air density on flight costs, as while wind results in savings or increases in the cost of transport depending on whether the bird is flying with or against it (*cf*.^20,21^), reductions in air density due to higher temperatures only result in increased flight costs (per unit time and per unit distance). This may therefore become more pertinent with increasing summer temperatures^22,23^. Temperature can also affect flight costs and capacities through its influence on other parameters, such as the development of thermal updrafts^24^. Red-tailed tropicbirds do extract energy from thermals over the ocean, however, a previous study found this was equally likely in summer and winter^25^, as the difference in temperature between the air and the sea remained constant across seasons.

Air density also varies in space, due to regional weather systems and global temperature gradients. We can estimate the corresponding geographical variation in flight power by extracting the air density across the red-tailed tropicbird distribution (Fig. 1). Within this range, the average power requirements are predicted to vary by 4.6 W in the austral summer (December, January, February) and 5.6 W in winter (June, July, August) (using the proportionality given in Bishop *et al*. ^6^). This is equivalent to 10-12% of the median power requirements across their range (47.4 W). The cumulative cost of flying in low density air could be substantial given the percentage difference in costs per second. This demonstrates that spatial variation in air density can produce substantial differences in flight power at sea-level, even when density values are strongly averaged (here they were taken across three months and averaged over 20 years for each season). We therefore predict that spatial variation in flight costs will result in large-scale differences in habitat quality.

Indeed, converting air density to density altitude (the altitude at which a particular air density occurs under standard atmospheric conditions, STAR methods), provides insight into the true extent that air density causes latitudinal changes in the flight environment (Fig. 2). On average, birds flying at sea level in the tropics experience air densities equivalent to 700 m above those operating at mid-latitudes (with substantial variation around the means, Fig. 2). Furthermore, birds operating at the highest latitudes in winter experience air densities equivalent to ∼ 1.5 km below sea level in standard atmospheric conditions. Therefore, whilst the expression “at sea-level” represents a useful baseline for height measurements, the same is not true for flight conditions, as latitudinal differences in temperature effectively create invisible topography at sea level, with a range of > 2 km when looking at long term averages (we used seasonal averages over two decades).

**Figure 2.**
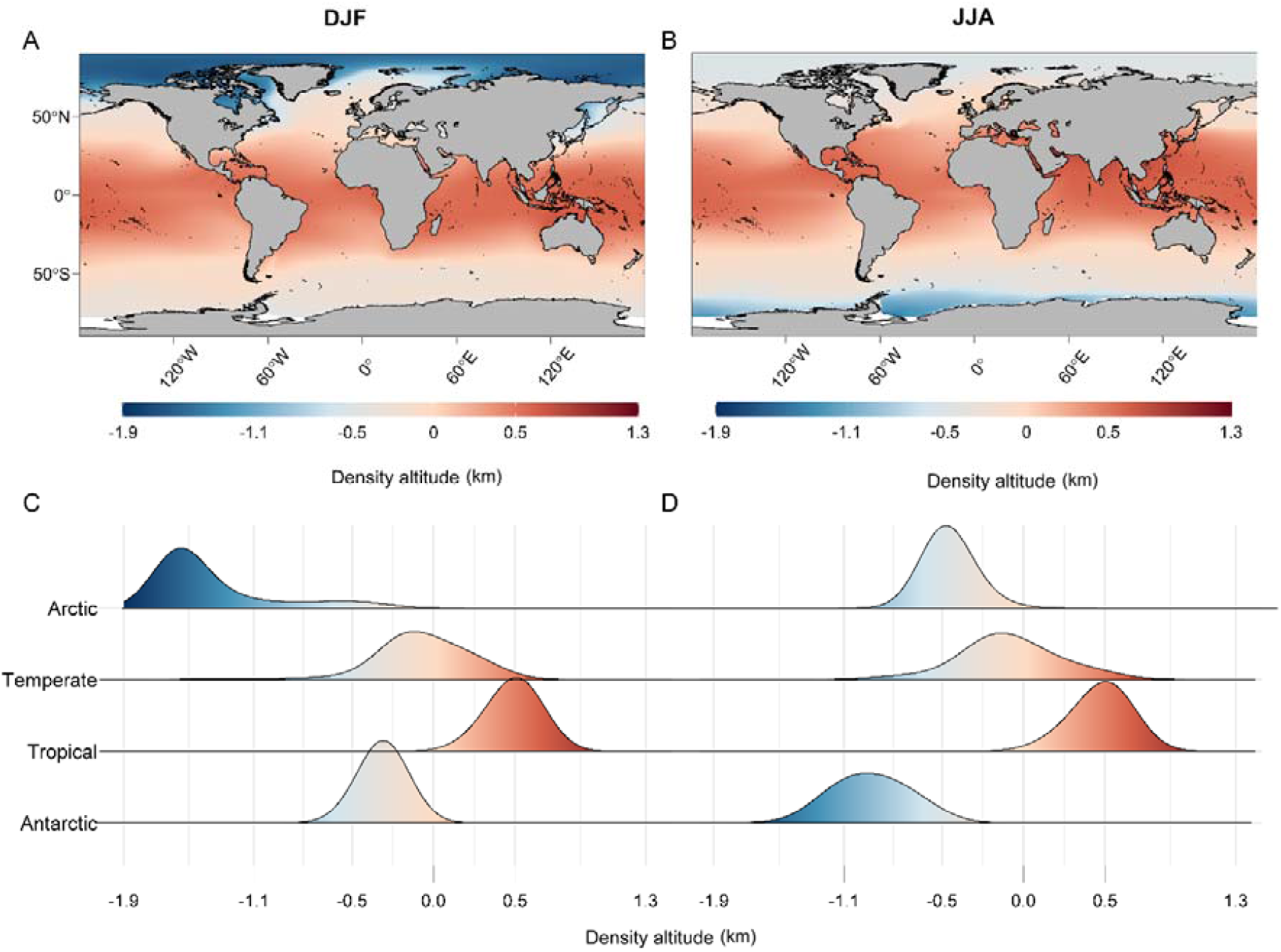
The effective altitudes, hereafter “density altitudes” (top two panels), experienced by any bird flying at sea level in two seasons (December, January and February, left panel, and June, July and August, right panel). Density altitude is the altitude corresponding to a given air density when compared to the reference density at sea level for the international standard atmosphere (ISA) (1.255 kg m^-3^) (see SI). The relative frequencies of the density altitudes are shown below the maps for each latitudinal zone. Tropical and temperate latitudes have modal peaks around 700 m and -200 m respectively.

This leads to the broader question of whether geographical trends in air density could affect the distribution of flying animals according to their morphology and flight style, being particularly pertinent for large-bodied fliers that experience disproportionately high costs of flapping flight (*cf*. Fig. 1). For instance, it is notable that the auks, which have the highest wing loading of all flying birds, occur in areas with higher air density. Their association with colder environments has been linked to their competitive ability to pursue prey^26,27^, but this would also have advantages in terms of lower flight costs, as well as an improved ability to dissipate heat during flight (*cf*.^28^). Overall, therefore, while temperature-driven changes in density have not previously been considered an important factor influencing flight costs, large-scale changes in temperatures may have affected the biogeography of flight morphologies, which will be important to understand against the backdrop of changing temperature regimes.

## Acknowledgements

This work was funded by a European Research Council starter grant (715874 to ELCS), under the European Union’s Horizon 2020 Research and Innovation program. We are grateful to the staff of Round Island for their support and the National Parks and Conservation Service, Government of Mauritius, who provided permission to access Round Island.

## Author contributions

ELCS conceived and supervised the project. Data were collected by AF, ELCS and NCC. Analyses were conducted by BG, EL and KK. The manuscript was written by ELCS, BG and EL with input from all authors. All authors gave approval for publication and agree to be held accountable for the work performed therein.

## Data availability statement

Data and code used for the analyses of this manuscript are available from the Zenodo Repository https://zenodo.org/records/13348839.

## METHODS

Breeding red-tailed tropicbirds were tagged on Round Island, Mauritius (19.8486° S, 57.7885° E), where breeding activity peaks between August and October (when > 25% of nest sites are occupied). The lowest breeding activity occurs between January and April (< 5% of nests are occupied (unpublished results^29,30^). During chick rearing, birds were equipped with a Daily Diary tag (Wildbyte Technologies, Swansea University, UK) and a GPS logger (GiPSy 5, Technosmart Europe, Guidonia-Montecelio, Italy) as detailed in Garde *et al*. ^25^. The Daily Diary recorded acceleration and magnetic field strength, each in 3 axes and at 40 and 13 Hz respectively. Barometric pressure and temperature were logged at 4 Hz, and GPS location was recorded once per minute. Both loggers were placed in a zip-lock bag and fixed to the back feathers using Tesa tape^31^. The loggers, housing and tape weighed 27.7 g, representing < 3% of the average body mass (mean body mass for tagged birds was 826 g), and 4.3% of the lowest body mass recorded during this study (650 g). Birds were weighed and photographed to quantify wing area and loading following Pennycuick ^10^. Ethical permission was granted by Swansea University AWERB, permit 040118/39.

Wind speed, direction, atmospheric pressure, relative humidity and temperature were recorded every 5 minutes by a portable weather station (Kestrel 5500L, Kestrel instruments, USA) mounted on a 5 m pole stationed at the highest point of Round Island (280 m ASL). Wind records for 7 flights were interrupted due to battery failure (between 9^th^ and 20^th^ February 2018). Here wind data were replaced by hourly wind records from Sir Seewoosagur Ramgoolam International Airport in Mauritius (downloaded from http://www.wunderground.com). Temperature, pressure, wind speed, wind direction and relative humidity were synchronised with the GPS data and linearly interpolated to 1-minute intervals.

### Flight metrics

Flight was evident as periods with variable altitude and was categorised as flapping or non-flapping flight using a simple acceleration threshold^25^. Specifically, the Vectorial sum of the Dynamic Body Acceleration (VeDBA) was calculated according to Wilson et al. ^32^, using smoothed raw acceleration values over two seconds to derive the gravitational component. VeDBA values were themselves smoothed over two seconds (sVeDBA) to produce a metric that varied between high and low levels of activity. A threshold of ≥ 0.4 *g* was used to identify flapping flight, which distinguished between flight types across individuals and seasons. The identification of flapping and gliding was undertaken in R Studio version 4.0.0^33^.

Wingbeat frequency was used as a proxy for flight effort rather than DBA^34,35^, as we identified a difference in the stability of the accelerometer attachment between the two seasons, which led to consistent differences in the amplitude of the acceleration signal^15^. Wingbeat frequency was robust to small changes in logger attachment and has been used to estimate work rate in a range of studies (e.g.^34,35^, though see^36^). Individual wingbeats were identified from peaks in the dynamic heave acceleration, smoothed over 3 events (0.075 s) following Krishnan *et al*. ^36^. In brief, peaks were identified as the highest values in the heave acceleration that occurred within five points of the positive to negative turning point in the rate of change of the heave acceleration. Each segment between successive peaks was counted as a wingbeat cycle, and the duration was used to calculate wingbeat frequency. Frequencies < 4 Hz were considered erroneous for birds of this body mass ^10^ (potentially resulting from manoeuvring or peak misidentification) and were therefore removed.

Flight altitude above sea level was calculated using the barometric pressure recorded by the Daily Diary (4 Hz), corrected for daily changes in sea-level pressure (taken from https://earth.nullschool.net/). Pressure values from the tags were smoothed over 2 s and the rate of change of altitude (*V*_z_) was calculated over 1 second intervals. The air density at flight altitude was estimated using the ideal gas law, using pressure measured by the Daily Diary and the temperature and relative humidity recorded by the weather station.

The bird’s groundspeed (*V*_g_) was taken as the haversine distance between GPS fixes divided by the time. The airspeed (*V*_a_) was estimated following Pennycuick ^10^, taking wind values from the anemometer. The headwind and crosswind components (HWC and CWC) were calculated as the two components of the wind acting on airspeed. A positive HWC indicates a headwind and a negative HWC corresponds to a tailwind.

Red-tailed tropicbirds are surface feeders ^37^ with regurgitates from birds on Round Island being mostly made up of flying fish. Prey pursuits were evident as rapid losses of altitude, temporary cessation of flapping flight and characteristic changes in the acceleration data indicative of manoeuvring (Fig. S2). These events were identified manually using custom-written animal movement analysis software DDMT (Wildbyte Technologies, http://wildbytetechnologies.com/software.html). We assumed that these represented most prey-encounters as there were no other obvious periods with bursts of acceleration and birds only ever dipped underwater for periods < 1.5 seconds, with these brief submersions usually occurring at the end of aerial pursuits.

### Statistical analysis

Seasonal differences in environmental and foraging parameters were tested as follows. Wilcoxon tests were used to test for differences in the body mass and wing area of tagged tropicbirds, as well as wind speed and temperature. Student’s t-tests were used to compare air density and chick age. Wind speed, temperature and air density were averaged over all flights per day in these tests to avoid pseudo-replication. Simple LME models were used to test whether total distance covered, wingbeat frequency, prey encounter rate and foraging efficiency varied between the two seasons, with individual as a random factor.

Birds vary their airspeed in relation to the wind, increasing their airspeed in headwinds and reducing it in tailwinds^20^. We therefore expected that we would be able to use either airspeed or wind in our model of wingbeat frequency and ran an LME to assess how much variation in airspeed was explained by wind and air density (birds are also expected to increase their airspeed with decreasing air density) ^7,38^. The model included the fixed effects of headwind and crosswind components, as well as wind strength, to test whether the estimated head and crosswind components captured all the variation due to changing wind conditions. Flight altitude was also included in the model to account for potential changes in currency (and associated speed selection) between relatively high and low altitude flight. This model was constrained to periods of level flapping flight (taken as periods where -0.2 < *V*_z_ < 0.2 and smoothed VeDBA was > 0.3 *g*), to control for changes in airspeed that occur in relation to climbing and descending flight ^39^. Individual bird ID was included as a random factor nested within foraging trip to account for unmeasured differences between days and individuals. We tested for collinearity by calculating the Variance Inflation Factors (VIF) of every fixed effect using the package “performance” ^40^. An autocorrelation structure of order 1 was also integrated to the model to account for the high level of autocorrelation in the GPS data (lag = 40).

Finally, we used an LME model to examine the factors causing the seasonal difference in wingbeat frequency. Here, we averaged values of airspeed, air density and flight altitude between successive prey pursuits to distinguish the effects of increasing body mass due to foraging success, from the effects of the physical environment. Only foraging trips with 5-15 pursuits and inter-pursuit intervals > 1 minute were included, resulting in 345 segments recorded over 48 trips from 41 birds. One outlier segment with a mean altitude > 250 m was excluded, as all other segments had a mean altitude < 150 m. The number of pursuits was included as a fixed effect in interaction with season, to identify any seasonal differences in foraging success. The presence of airspeed in the model accounted for the expected effects of the wind on the wingbeat frequency. Air density and altitude were both included in the model to enable us to distinguish between the effects of changing flight altitude and seasonally changing temperatures. “Individual” was used as random factor nested within “foraging trip”. Statistical analyses were carried out using R Studio, using the packages nlme (Pinheiro et al. ^41^, version 3.1-151) and MuMIn (Barton and Barton ^42^, version 1.43.17).

### Estimation of density altitude

The following equations describe temperature, pressure and density of the air in the international standard atmosphere (ISA) within the troposphere, which extends up to 11 km altitude:

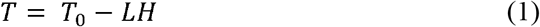

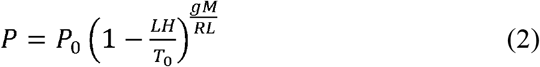

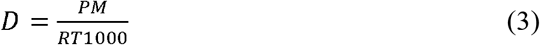

where *T* = ISA temperature in deg K

*P* = ISA pressure in Pa

*D* = ISA density in kg/m^3^

*H* = ISA geopotential altitude in km

These can be rearranged to express geopotential altitude as a function of density:

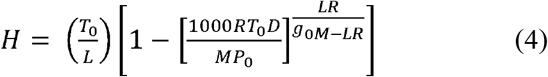

This can be simplified to the following when ISA constants are used:

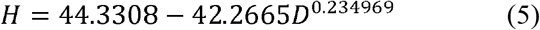

where H = geopotential altitude in kilometres.

