## Supplementory information for "Latitudinal gradients in air density create “invisible topography” at sea level affecting animal flight costs"

### **Supplementary Information**

***Table S1:*** *Weather conditions and flight performance in the Austral summer (February-March) and winter (September-October). Values are the means ± SD. The relationships between season, wind direction and outward heading were tested using Watson-Wheeler tests, whereas the relationship between all other variables was tested using linear mixed effect models (LME ^23^). Significance is indicated according to p-value: p < 0.001 (***), p < 0.01 (**), p < 0.05 (*).*

| **Variable** | **Summer** | **Winter** | **Significance** |
| --- | --- | --- | --- |
| Air density (kg m^-3^) | 1.16 ± 0.01 | 1.19 ± 0.01 | ** |
| Wind direction (degrees) | 81.9 ± 0.9 | 89.2 ± 0.2 | ** |
| Wind speed (m.s^-1^) | 5.0 ± 3.5 | 6.3 ± 2.2 | NS |
| Trip distance (km) | 83.5 ± 58.1 | 111.5 ± 105.4 | NS |
| Trip duration (hours) | 5.1 ± 7.8 | 5.6 ± 8.2 | NS |
| Outward heading (degrees) | 158.6 ± 1.0 | 154.5 ± 1.1 | NS |
| Airspeed (m s^-1^) | 9.1 ± 2.4 | 9.8 ± 2.2 | NS |
| Mean flight altitude (m) | 69.8 ± 23.3 | 55.5 ± 16.5 | ** |
| Passive flight (% of time) | 12.0 ± 8.7 | 13.4 ± 6.9 | NS |
| Wingbeat frequency (Hz) | 4.0 ± 0.1 | 3.9 ± 0.1 | ** |
| Number of pursuits per trip | 8.5 ± 6.4 | 8.0 ± 6.6 | NS |
| Pursuit distance from colony (km) | 21.8 ± 16.6 | 27.2 ± 26.8 | NS |
| Foraging efficiency  (pursuits/ wingbeats per trip * 10,000) | 2.8 ± 1.7 | 2.2 ± 1.4 | NS |

***Table S2.*** *Effect of the physical environment on airspeed selection.* *Output of the LME model showing the effect of air density, altitude, headwind component (HWC), crosswind component (CWC) and wind speed on airspeed. R²_m_ = 0.18, R²_c_ = 0.25.*

|  | **Estimate**  **(raw)** | **Estimate**  **(scaled)** | **Std. Error** | **t-value** | **p** | **VIF**  **(< 5)** |
| --- | --- | --- | --- | --- | --- | --- |
| (Intercept) | 49.522 | 11.740 | 12.669 | 3.909 | < 0.001 | NA |
| Air density (kg m^-3^) | -35.305 | -0.579 | 10.655 | -3.313 | 0.001 | 2.09 |
| Altitude (m) | 0.019 | 0.579 | 0.002 | 8.685 | < 0.001 | 2.08 |
| HWC (m s^-1^) | 0.211 | 0.818 | 0.017 | 12.055 | < 0.001 | 1.52 |
| CWC (m s^-1^) | -0.036 | -0.079 | 0.023 | -1.604 | 0.109 | 1.35 |
| Wind speed (m s^-1^) | 0.376 | 1.033 | 0.038 | 9.886 | < 0.001 | 1.41 |

***Table S3****. Wingbeat frequency of red-tailed tropicbirds.* *Output of the LME model showing variation in wingbeat frequency as a function of air density, airspeed, flight altitude and pursuit number in the low and high season, as well as the effect of airspeed on the slope between air density and wingbeat frequency.* *R²_m_ = 0.13, R²_c_ = 0.77.*

|  | **Estimate (raw)** | **Estimate (scaled)** | **Std. Error** | **t-value** | **p** |
| --- | --- | --- | --- | --- | --- |
| (Intercept) | 10.881 | 4.420 | 0.029 | 149.943 | < 0.001 |
| Air density (kg m^-3^) | -5.392 | -0.084 | 0.029 | -2.913 | 0.004 |
| Airspeed (True, m s^-1^) | -0.312 | 0.002 | 0.002 | 1.214 | 0.226 |
| Altitude (m) | -0.003 | -0.061 | 0.008 | -7.257 | < 0.001 |
| Air density: Airspeed | 0.266 | 0.004 | 0.002 | 2.463 | 0.014 |
| Season:Pursuits (Low) | 0.008 | 0.031 | 0.007 | 4.200 | < 0.001 |
| Season:Pursuits (High) | 0.010 | 0.039 | 0.006 | 6.112 | < 0.001 |
| Season*Pursuits | 0.002 | 0.007 | 0.009 | 0.792 | 0.428 |

***Figure S1.*** *Seasonal variation in average monthly wind speed, temperature and air density in Mauritius as averaged between 2015 and 2019 (with values taken from ERA5), compared to the monthly variation in the proportion of nest sites occupied by tropicbirds on Round Island in 2017 ^24^. The shaded regions correspond to the summer (light grey) and winter (dark grey) breeding seasons.*

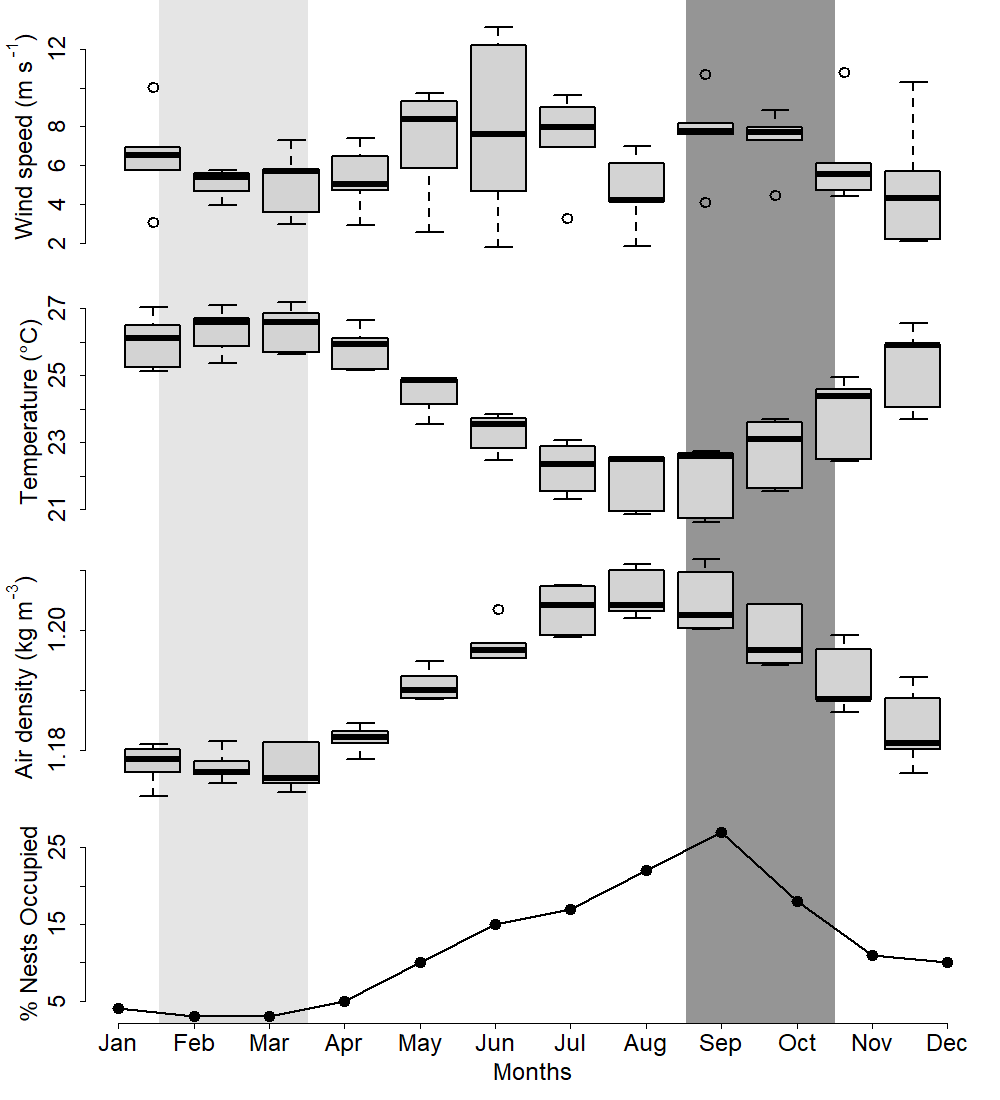

***Figure S2.*** *Red-tailed tropicbird movement data.* *Prey pursuits were typified by the following parameters from top to bottom: loss of altitude, rapid increase in vertical velocity (*V*_z_) (negative values indicate descending flight), and a temporary switch from flapping to passive flight, as evident in the heave (dorsoventral) acceleration values.*

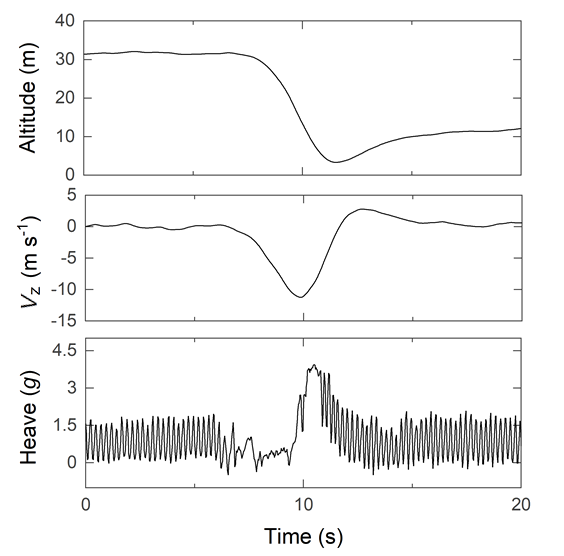
